# Muscle synergies are associated with intermuscular coherence in an isometric upper limb task

**DOI:** 10.1101/843797

**Authors:** Pablo Ortega-Auriol, Winston D Byblow, Angus JC McMorland

## Abstract

To elucidate the underlying physiological mechanism of muscle synergies, we investigated the functional corticomuscular and intermuscular binding during an isometric upper limb task in 14 healthy participants. Cortical activity was recorded using 32-channel encephalography (EEG) and muscle activity using 16-channel electromyography (EMG). Using non-negative matrix factorization (NMF), we calculated muscle synergies from two different tasks. A preliminary multidirectional task was used to identify synergy preferred directions. A subsequent coherence task, consisting of generating forces isometrically in the synergy PDs, was used to assess the functional connectivity properties of synergies. Functional connectivity was estimated using corticomuscular coherence (CMC) and intermuscular coherence (IMC). Overall, we were able to extract four different synergies from the multidirectional task. A significant alpha band IMC was present consistently in all extracted synergies. Moreover, alpha band IMC was higher between muscles with higher weights within a synergy. In contrast, no significant CMC was found between the motor cortex area and synergy muscles. In addition, there is a relationship between a synergy muscle weight and the level of IMC. Our findings suggest the existence of a consistent shared input between muscles of each synergy. Finally, the existence of a shared input onto synergistic muscles within a synergy supports the idea of neurally-derived muscle synergies that build human movement.

## Introduction

Dimensionality reduction applied to EMG signals from multiple muscles shows the presence of lower-dimensional structures, which we call muscle synergies that can explain the behaviour of the full set of muscles measured. There are two key unanswered questions relating to these muscle synergies that we will address in this study. Firstly, do muscle synergies represent a deliberate neurophysiological control strategy (Bizzi and Cheung 2013; McMorland et al. 2015) or are they merely an artefact of the task requirements and mathematical derivation (Kutch and Valero-Cuevas 2012)? Second, if muscle synergies arise from a control strategy, which neural structures are responsible for their emergence and modulation?

Evidence exists on both sides of the debate relating to whether synergies arise from neural structures. Animal (Tresch and Bizzi 1999; Bizzi et al. 2008; Hart and Giszter 2013) and computational (Neptune et al. 2009) models using electrical stimulation support a neural origin for muscle synergies. Similarly, human experiments are consistent with a neural origin of muscle synergies during natural movements (d’Avella and Bizzi 2005; Safavynia and Ting 2012), affecting learning rates (Berger et al. 2013; Sawers et al. 2015), for postural control (Weiss and Flanders 2004), when extracted from the frequency domain (Frere 2017), irrespective of muscle fatigue (Ortega-Auriol et al. 2018), and in the presence of CNS damage after stroke (Cheung et al. 2009; Berger et al. 2013). Conversely, evidence for muscle synergies as a result of purely mechanical constraints arise from computer simulations of the upper arm movement on a single plane (Inouye and Valero-Cuevas 2016) and cadaveric studies (Kutch and Valero-Cuevas 2012). While it seems possible that muscle synergies may arise from both neural and biomechanical mechanisms, there are still good reasons to determine which neural structures are implicated in their expression.

A viable theory about movement control must contain a neuroanatomical framework that is capable of discerning the origin of muscle synergies (McMorland et al. 2015). Movement control can be deconstructed into three main sources of drive: cortical activity, spinal activity through central pattern generators, and adjustment reflexes (Ivanenko et al. 2005, p. 200). A number of experimental approaches exist that can differentiate between the possible neural sites of origin of motor behaviours. Coherence, a signal frequency-based analysis, is capable of identifying common functional control sources across muscles during a task (Laine and Valero-Cuevas 2017). Coherence is a measure of correlation between two signals in a determined frequency band (Boonstra 2009). Cortico-muscular coherence (CMC) between brain (EEG) and muscle (EMG) activity, occurs around the beta band (15–30 Hz), suggestive of cortical control of movement (Gwin and Ferris 2012). On the other hand, intermuscular coherence (IMC), between EMG of different muscles, occurs around the ∼10 Hz or alpha band (Boonstra 2009) and is considered to reflect subcortical control (Boonstra 2009; Marchis et al. 2015). We can use coherence to identify the neurophysiological sources of drive that generate muscle synergies.

On a functional level, CMC and IMC are measures that can provide essential insight into the underlying coordination of a motor task, differentiate pathways converging onto spinal motor neurons, and extract shared information across different scattered muscles (Boonstra 2013). Synchronisation of oscillatory activity across the neuromuscular system as measured by coherence reflects the necessary functional coupling to control movement. Factors such as cortical spectral power (Kristeva et al. 2007) and sensory feedback (Fisher et al. 2002) can modulate IMC levels. Synchronisation within the beta band during precise, steady force outputs of hand muscles have been proposed to be indicative of functional binding between the primary motor cortex and effector muscles (Kilner et al. 2000; Boonstra 2009; Danna-Dos Santos et al. 2010). Alpha band synchronisation across distinct muscles or even bilateral muscles, as IMC, suggests a common input from subcortical structures (Conway et al. 1995; Baker et al. 2003) and may specifically reflect the involvement of the reticulospinal pathway (Grosse and Brown 2003)

Coherent activity has been described across muscles of an individual synergy during postural responses (Danna-Dos-Santos et al. 2014) and a cycling task (Marchis et al. 2015). However, coherent activity (either beta or gamma bands) was found in both studies only within a single synergy out of the three or more extracted, suggesting cortical modulation. Two main concerns arise from the cited results: first, coherent activity was only present within a single synergy, implying a non-neural origin or lack of task functionality for the rest of the extracted synergies during a motor task; second, these studies did not include EEG recording, precluding the confirmation of a cortical origin of the other extracted synergies. At the same time, a functional task might not be an optimal paradigm to determine coherent activity within synergies. Instead, trials tuned to the preferred direction of a synergy will preferentially recruit that single synergy. The recruitment of a single synergy may lead to increase the robustness and coherence across trials by reducing the possible noise emerging from multiple synergies being recruited at the same time.

Our aim is to develop an approach able to provide insights on the nature of muscle synergies, either arising from a functional neural network or as a result of mechanical or mathematical constraints. If synergies arise from purely mechanical/mathematical concerns, we would not expect to see coherence between muscles. Conversely, if synergies truly arise from functional neural connections with a common source, muscles within a synergy should be coherent during task performance. Moreover, since the drive to muscles is excitatory, common information creating the structure of a synergy is disproportionately distributed to muscles with higher weights in a synergy, and therefore it is likely that these muscles specifically would exhibit higher coherence. We hypothesised that there would be coherent activity within synergies during the performance of an isometric contraction of the upper limb (UL). Specifically, we hypothesised that if a synergy originates from cortical modulation, some muscles, especially those with a higher weight within that synergy, would exhibit a significant level of CMC (∼20Hz band). Alternatively, if the spinal level or other subcortical circuitries modulate synergies, those muscles with a higher weight contribution should present a significant level of IMC (∼10Hz band coherence).

## Materials and Methods

### Participants

We recruited 14 right-handed volunteer participants (Table 1); participants were young and healthy without any pathology that affected the UL, spine or posture. Volunteers were excluded if they reported neck, shoulder or arm pain (> 2 in a 1-10 verbal scale) within the last three months. The University of Auckland Human Participants Ethics Committee approved the research protocol and methods of the study (reference number 022246), and informed consent was gained before participation in any procedure.

**Table 1.**
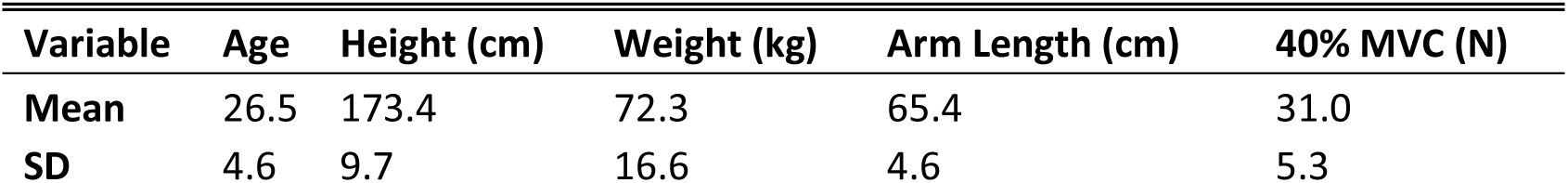
Participants characteristics.

### EEG and EMG

EEG signals were recorded from a 32 Ag/AgCL electrode EEG system (EasyCap; Brain Products GmbH, Germany). The electrodes were positioned according to the 10–20 system, referenced to the FCz channel, and offline to a common reference. Signals were acquired with BrainVision Recorder software (Brain Products GmbH, Germany) at 5 kHz.

Surface EMG signals were recorded from 16 single differential channels and sampled at 2 kHz using a Trigno System, (Delsys Inc., United States). EMG activity was recorded from muscles of the participant’s dominant UL: superior (ST) and middle trapezius (MT), infraspinatus (Inf), teres minor (TM), serratus anterior (SA), anterior (ADel), middle (MDel), and posterior deltoid (PDel), pectoralis major(PM, clavicular fibres), short (BS) and long (BL) heads of biceps brachii, long (TL) and lateral (Tlat) heads of triceps brachii, brachioradialis (Braq), extensor carpi radialis (ECR), and flexor carpi radialis (FCR). These muscles were chosen on the basis of their force capability and likely contribution to the required task, muscle characteristics which are essential for accurate reconstruction of synergies (Steele et al. 2013). Electrodes were positioned according to Seniam and Cram’s guidelines (Hermens et al. 1999; Criswell 2010). Each participant’s skin was prepared with an abrasive gel. Signals were recorded via a custom software interface.

### Tasks and Protocol

The experiment took place on a single session, where participants performed three tasks: maximal voluntary force (MVF), multidirectional trials, and synergy tuned trials. Part of the experimental protocol has been described previously (Ortega-Auriol et al. 2018). While seated the participants exerted 40% of the MVF over a handle instrumented with a force transducer (Figure 1a) located at 40% of the arm length (acromion – 3^rd^ metacarpal head) in front of the shoulder. Forces were recorded at 120 Hz with an instrumented handle with a 6-axis force transducer (Omega160, ATI Industrial Automation, United States). Signals were recorded by a custom software interface. Force level and direction were feedback to the participant by virtual reality feedback (VRF). The VRF consisted of a 3D force space, where a movable sphere had to be matched to a specific target (Figure 1d). For both tasks, the required force to match the target was equivalent to the participant’s 40% MVF defined as the maximum force from three trials of external shoulder rotation.

**Figure 1.**
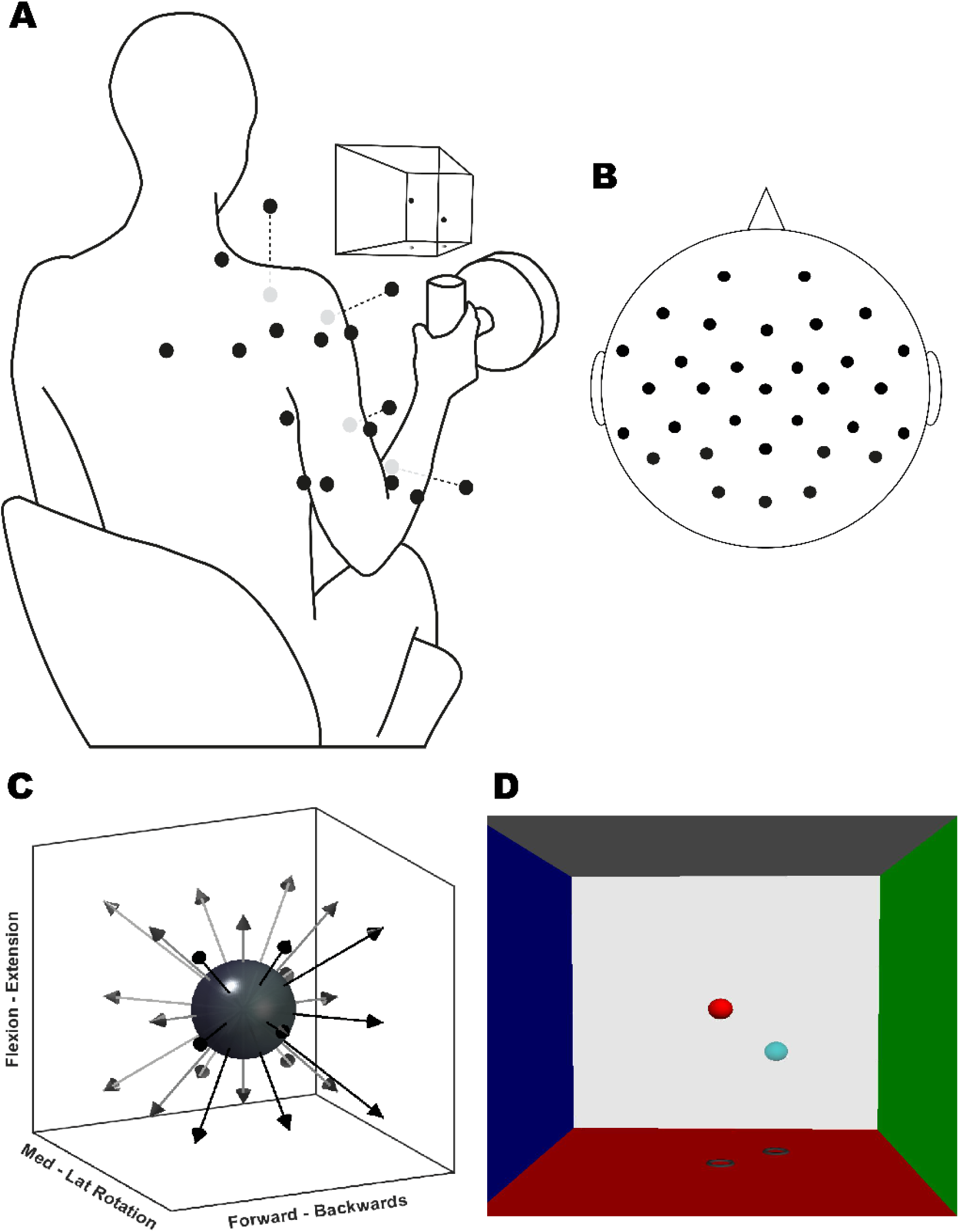
Experimental setup. **(A)** Illustration of EMG sensor placement (black dots, grey dots are located ventrally), VRF, and instrumented handle. **(B)** 10–20 EEG setup schematic. **(C)** Representation of target directions of the multidirectional task. **(D)** Screenshot of the VR feedback displayed on the screen. Each VRF wall is located 100 N away from the centre.

The multidirectional task consisted of isometric contractions to match targets in 26 different directions evenly distributed around a sphere (Figure 1c). A trial was considered valid after four seconds matching the target within a range of ±5 N. Signals were recorded from the end of the previous trial until four seconds had elapsed of continuous target match. From the multidirectional task EMG processing, three types of variables were identified for each participant: (i) the coefficients of muscles synergies which determine synergy structure, (ii) the significant number of synergies able to reconstruct the original EMG data to a threshold level of variance accounted for (VAF), and (iii) the spatial tuning of each significant synergy or synergy preferred direction (PD) determined by the level of activation of the synergy in the different directions.

The synergy tuned trials consisted of a sustained isometric contraction towards a specific SPD again by matching a specific target. For a trial to be valid, participants had to match a target for four consecutive seconds. Once this time was reached, participants released the handle before the start of the next trial. To corroborate synergy structure, synergies were extracted again from the concatenated trials of each PD. The participants performed 50 trials in each PD to calculate CMC and IMC. Also, participants performed 20 trials at 25, 50 and 75% of the distance in the plane between the neighbour synergies. Neighbour synergies were first defined as a pair of synergies with the lowest angle in a vector space. The number of neighbour synergies was defined as the number of significant synergies minus one. The order of the trials was randomised and self-paced; participants were able to rest between trials to avoid fatigue effects.

### Data Analysis

Synchronisation across devices, EMG and force acquisition, were performed with Python-based custom software (https://dragonflymessaging.org/applications.html, U. of Pittsburgh). Data analysis was performed in MATLAB 9.3 (MathWorks, United States) using custom made scripts and the FieldTrip toolbox (Oostenveld et al. 2011). A schematic representation of the workflow to process the data is shown in Figure 2.

**Figure 2.**
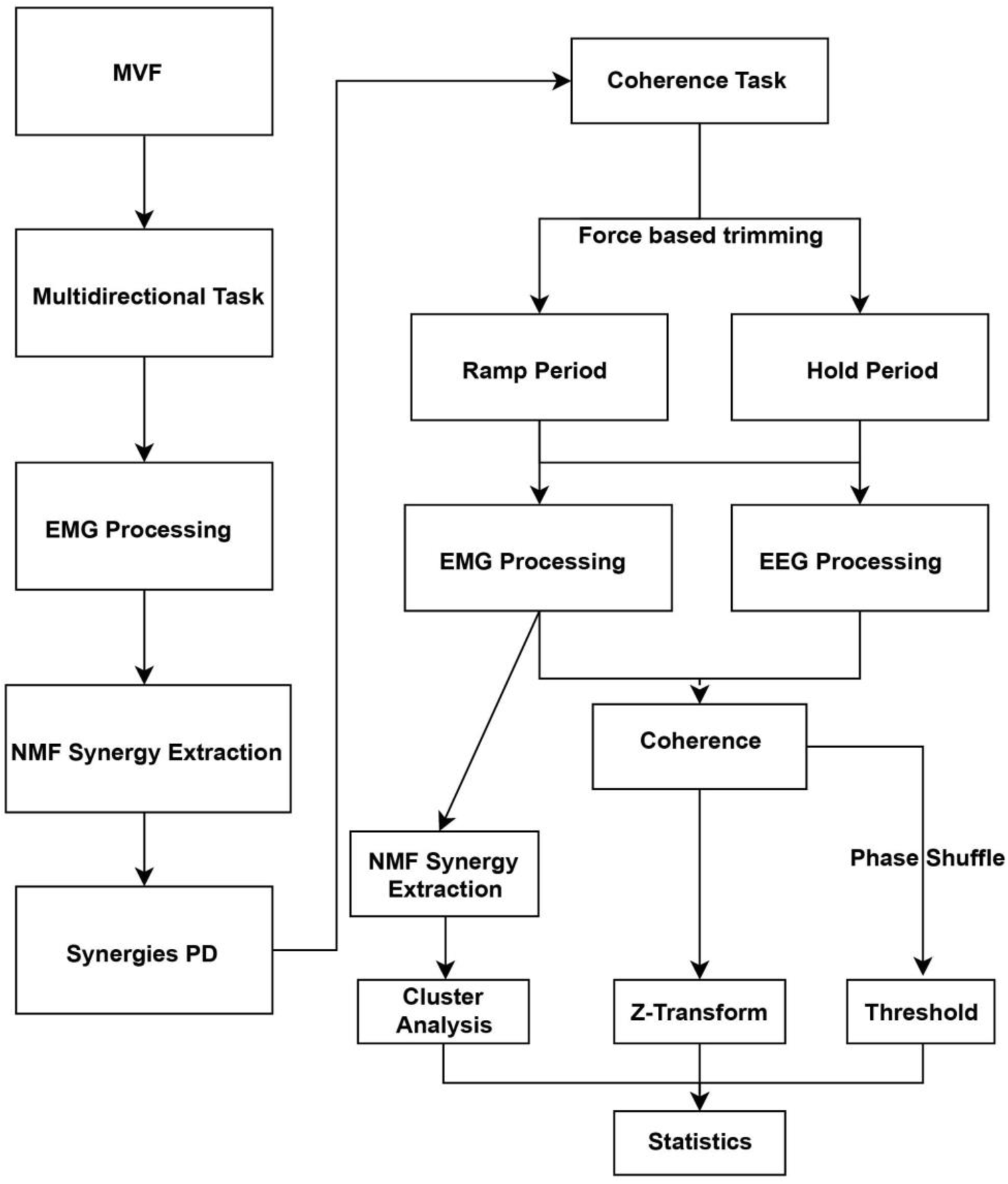
Data processing pipeline from MVF to statistical analysis.

#### EMG processing

The processing of multidirectional trials for synergy extraction is described in detail in our previous research (Ortega-Auriol et al. 2018). Briefly, EMG signals were: trimmed for the intermediate two seconds of the target match period, band-pass filtered (bidirectional Butterworth, 2^nd^ order, 5–400 Hz.), rebinned into 100 data points, demeaned, full-wave rectified, normalized to maximum activation during each trial across all muscles, converted to unit variance and low pass filtered again to obtain an envelope (Butterworth, 2^nd^ order, 5 Hz).

#### Synergy extraction

Non-negative matrix factorisation (Lee and Seung 1999) was applied to the processed concatenated EMG signals from the multidirectional task. NMF can be modelled as *D* = *W* * *C* + *ϵ*, where D is the original data set, W the synergy structure and C the activation coefficients, and *ϵ* is the unexplained variance not explained by the synergies. NMF was implemented using the multiplicative rule (Berry et al. 2007). The final solution was the result of 20 consecutive iterations with a difference of EMG reconstruction error smaller than 0.01%among them. To determine a significant number of synergies, first, the algorithm iterated from one until the number of muscles minus one. Secondly, we used the VAF metric (Cheung et al. 2005) to determine the number of synergies that achieved the best reconstruction of the original data. VAF was applied as a global (quality of original dataset reconstruction) and local criteria (quality of individual muscles signals reconstruction). A significant number of synergies was determined when global VAF ≥ 90% and local VAF ≥ 80%.

#### Synergy preferred directions

PDs were calculated for each extracted synergy. PD was defined as the average of all directional unit vectors of the multidirectional trials, scaled by the activation coefficient of the correspondent extracted synergies.

#### Coherence task trials pre-processing

EEG and EMG data from coherence trials were band-pass filtered (bidirectional Butterworth, second-order, 5–100 Hz.), demeaned, EMG data was rectified via Hilbert Transform, and each trial was split into a ramp and a hold phase. The ramp phase was defined as the time window from the initial movement of the VRF sphere until the inflexion point (knee) of the force trace. The hold phase was defined as the four seconds following the end of the ramp phase. To split the trials, force data were low pass filtered (Butterworth, second-order, 5 Hz), and the inflexion point of the force traces was calculated by a custom algorithm and corrected through visual inspection if necessary. EEG signal impedance was checked at the beginning of the session through the implementation and after the multidirectional trial. Impedance levels were checked between tasks and kept under 15 Ohms through the experiment. EEG data was downsampled to 2 kHz to match EMG sampling frequency and optimise processing times. An independent component analysis (ICA) was applied to remove electrooculographic artefacts; the component that visually presented EOG artifacts and was spatially located on the ocular region was subtracted from EEG data reconstruction.

#### Cluster analysis

To group similar synergies across participants, a cluster analysis was applied to the pooled synergy structures of all participants, derived from EMG data collected in the coherence task. Cluster analysis was applied using a k-medoids algorithm (Park and Jun 2009) with a cosine function as the cluster distance metric, and applying the Silhouette index (Kaufman and Rousseeuw 1990) to determine the number of clusters within the pooled synergies. A mean synergy, and the dominant muscles within each mean synergy, were calculated for each cluster. Dominant muscles were determined as those with highest weights contributions, with weights greater than the averaged mean weight plus one standard deviation within each synergy structure.

#### Coherence Calculation

Before calculating CMC and IMC, we concatenated all the trials in a single PD direction, for each of the PDs we had calculated from the multidirectional task. From each set of concatenated trials, we extracted a single synergy. We checked the resulting synergies for consistency with the calculated synergies from the multidirectional task. Structures from the re-calculated four synergies were not different from those extracted from the multidirectional task. Then, we calculated IMC and CMC from the 50 concatenated trials for each PD separately. We calculated and compared IMC between three different muscle groups: (A) high–all calculation, representing the average IMC of the three highest contributors muscles with all other muscles within a single synergy; (B) high–high being the average IMC only between the three highest contributors within a synergy; and (C) high–low displaying the average IMC between three highest and the three lowest contributors within each synergy.

EMG and EEG signals were transformed into the frequency domain. A fast Fourier Transformation (FFT) was applied using a multitaper for both ramp and hold phases independently. FFT was applied to the bandwidth between 3–50 Hz using three tapers. FFT results consisted of 24 frequency bins between 3–50 Hz with steps of 2 Hz. To narrow the scope of CMC calculations, we only considered channels on and near the motor cortex area: FC5, FC1, Fz, Cz, C3, T7, CP5, CP1. CMC and IMC were calculated for the available combinations between EEG – EMG and EMG – EMG channels. From the subset of analysed EEG channels, the one with the highest average CMC value was used for further analysis. Similarly, for IMC, the dominant muscles from each synergy were considered for the CMC statistical analysis of the data. In other words, IMC was the average value of the dominant muscles against all other muscles within a single synergy (the high–all data). Raw coherence values were normalised by applying a z-transformation (Baker et al. 2003; Reyes et al. 2017) applying equation 1:

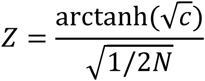

where c is the raw coherence value, and N the number of tapers used in the coherence calculation. From the individual estimates of coherence, pooled CMC and pooled IMC were calculated to produce a single global estimate of correlated CMC or IMC (Amjad et al. 1997). Pooled coherence was estimated using equation 2:

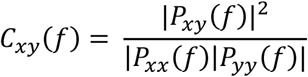

To define a significance level for CMC or IMC, a threshold was calculated based on coherence analysis of a surrogate time series derived from the original EMG and EEG data. Once the original data were transformed into the frequency domain, the phase components as the imaginary parts of the resultant complex number were independently shuffled. The shuffling was iterated 50 times among trial repetitions, channels, and participants. Then, coherence was calculated as described previously. This procedure allows the conservation of the power spectrum original amplitude structure of the signal while only shifting the signal phase, uncorrelating the signals in the time and frequency domain (Faes et al. 2004; Marchis et al. 2015). CMC and IMC threshold significance was established as above the 95th percentile of the resultant by-chance coherence distribution. To test our hypothesis, we compared the average CMC and IMC values above the threshold of high–high weight muscles against high–low weight muscles. A higher degree of coherence between muscles with a high–high weight would support a neural origin of synergies.

Finally, we compared the average IMC within different groups of muscles within each synergy (high–all, high–high, and high–low, Figure 5) using Friedman’s ANOVA between groups, this analysis was constrained within the relevant frequency band (7–16 Hz, Figure 4). Data distributions were first checked for normality by using a Kolmogorov-Smirnov Test (k - test), assessing the skewness value, and visually inspecting a normality plot.

**Figure 3.**
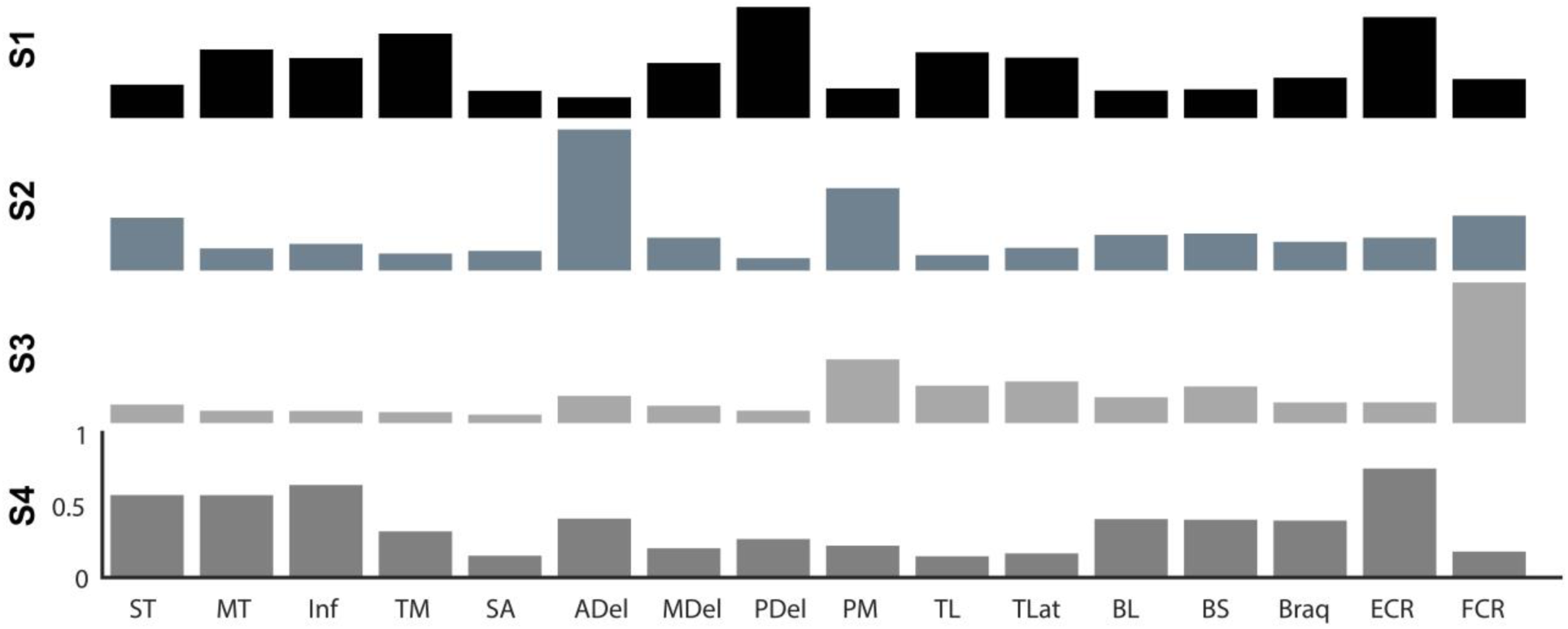
Synergy structures of each cluster (S1 – S4). Each bar represents the normalized muscle weight within each synergy. From top to bottom: S1 extension synergy (black), S2 flexion synergy (slate grey), S3 adduction synergy (light grey), and S4 external rotation synergy (grey).

**Figure 4.**
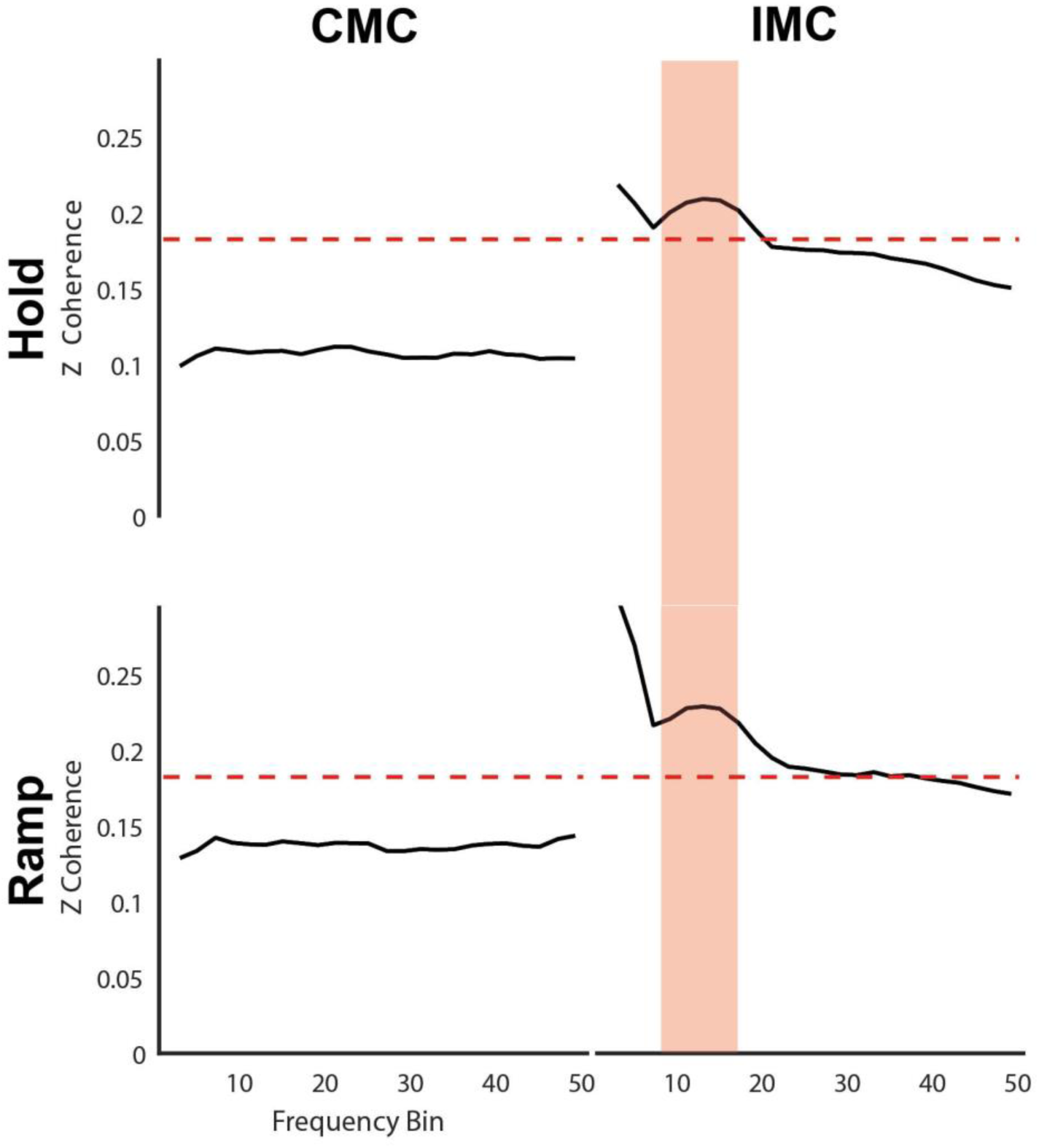
Mean Z-transformed CMC and high–all IMC across all clusters of the hold and ramp phases. We found a non-significant level of CMC in both phases, and a significant level of IMC around the alpha band in both phases. Dashed red lines show the significance threshold. The IMC shaded area represents the relevant frequency band analysed on figure 5.

**Figure 5.**
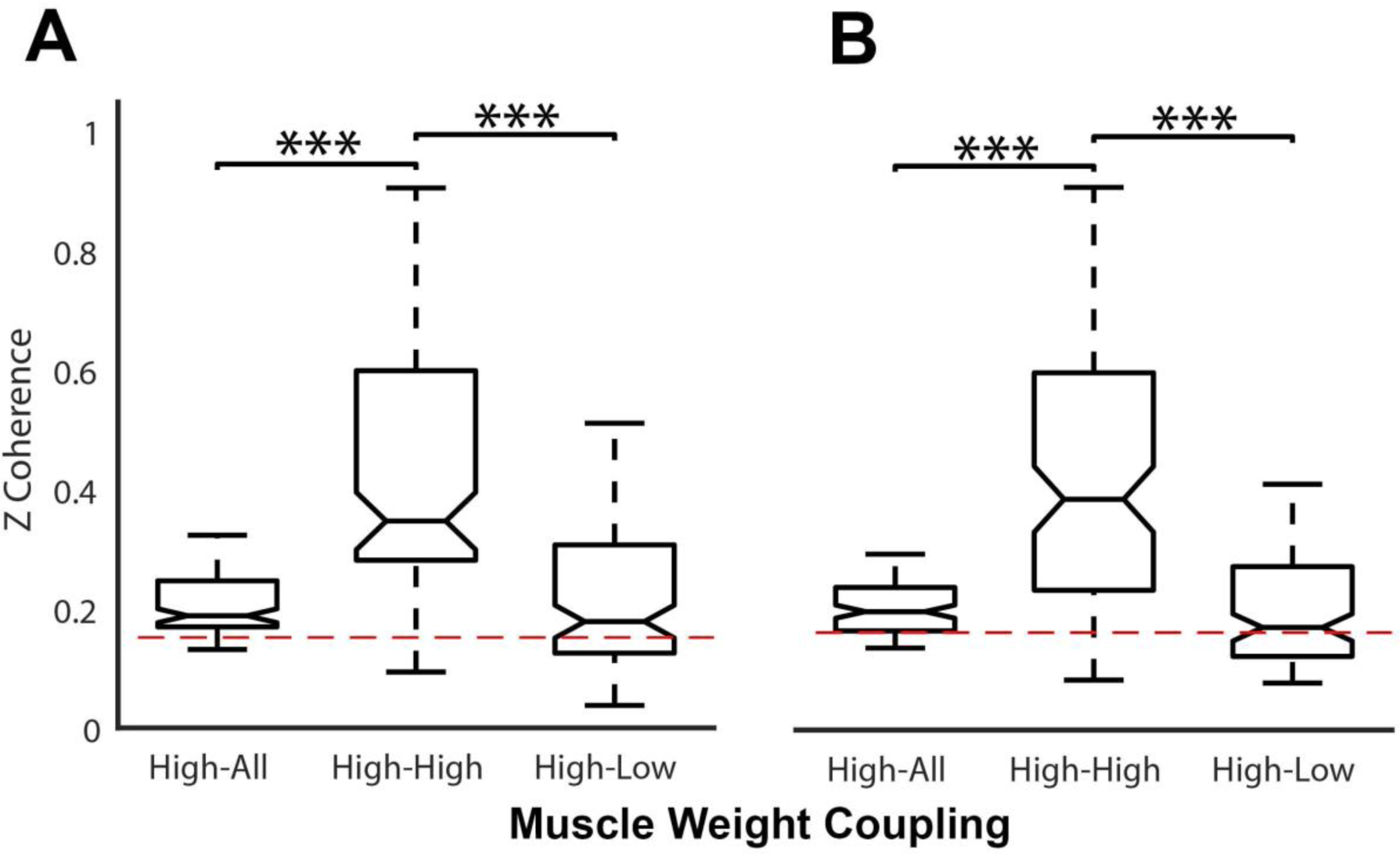
Average IMC values within the relevant frequency band across clusters comparing the different IMC calculations of (A) hold and (B) ramp phases. Dashed red lines show the significance threshold.

## Results

All participants were able to complete the multidirectional and coherence trials on the requested directions and repetitions. In average, from multidirectional trials, 4.2 (SD 0.6) synergies were sufficient to reconstruct the original EMG data set. The average MVF was 77.6 N (SD 12.2), requiring the exertion on average of 31 N (SD 5.3) as 40% of the MVF (Table 1).

### Muscle Synergies

Four muscle synergies were identified (Figure 3). S1 can be interpreted as a shoulder extensor synergy with the involvement of PDel, TM and a minor contribution from triceps muscles. S2 is a shoulder flexor synergy with contributions from the ADel and PM muscles. S3 performs adduction and internal rotation synergy with a higher weight of the FCR and PM. Finally, S4 is an external rotation synergy, with the contribution of the MT and ST as scapula stabilisers while the Inf muscle contributes to the rotation. This functional interpretation resembles an orthogonal distribution of the extracted synergies crossing the shoulder joint.

### CMC & IMC

CMC considered independently for each force PD, was below the by-chance threshold for both ramp and hold phases of the tasks (Figure 4). During the hold phase, synergies show a predominant high–all IMC coupling within the alpha band (∼10 Hz). Interestingly all four synergy clusters showed some extent of coherent activity (Table 2). For the ramp phase, we found similar results with a dominant coupling activity peak in the alpha band.

**Table 2.** Average IMC(sd) and frequency of the ramp and hold periods of each cluster.

| Hold Period |  |  |  |  |  |
| --- | --- | --- | --- | --- | --- |
|  | Cluster 1 | Cluster 2 | Cluster 3 | Cluster 4 | Clusters Mean |
| IMC (SD) | 0.22 (.02) | 0.19 (.01) | 0.18 (.00) | 0.2 (.01) | 0.20 |
| Frequency | 14.0 | 11.0 | 3.0 | 11.0 | 9.8 |
| Ramp Period |  |  |  |  |  |
| IMC (SD) | 0.24 (.05) | 0.23 (.03) | 0.2 (.01) | 0.21 (.03) | 0.22 |
| Frequency | 13 | 11 | 5 | 16 | 11.25 |

### High-low contributor

Distributions of IMC according to weight contribution, were not normally distributed [k-test p = 0.0002, skewness = 0.5] and low [k-test p = 0.002, skewness = 0.8] distributions were both significantly non-normal. Therefore, we checked the mean difference between the IMC distributions by applying a Friedman’s ANOVA. The IMC level was different across the three different IMC calculations for the hold (*x*^2^(2) = 97.0, *p* = 0.00008) and ramp (*x*^2^(2) = 75.7, *p* = 0.0007) phases. Wilcoxon tests were used to find individual group differences. For the hold phase, the high–high IMC was significantly different from high–low (z = 8.3, *p* = 0.0006) with a large effect size (ES) = 0.81 (Rosenthal 1986), and different from high–all IMC (z = 7.8, *p* = 0.0003) with a medium ES = 0.76; no significant difference was found between high–all and high–low IMC calculations. Similarly, for the ramp phase, the high– high IMC was significantly different from high–low (z = 8.1, *p* = 0.0005) with a medium ES = 0.78, and different from high–all IMC (z = 7.5, *p* = 0.0003) with a medium ES = 0.72, and no significant difference was found between high–all and high–low IMC calculations.

## Discussion

In order to determine the functional connectivity of muscles within a synergy, we quantified the correlations between EEG-EMG and EMG-EMG channels. We found three main results: first, a consistent, significant IMC level across synergies; second, muscles with a higher weight within a synergy showed a consistent higher IMC level; and third a lack of CMC on both ramp and hold periods. These findings suggest a subcortical involvement in the generation of muscle synergies for this isometric upper limb task.

### Corticomuscular coherence

We found no evidence of CMC in the present study. Several factors may explain this. Firstly, variable force output extinguishes CMC that can be seen in subsequent periods of constant force production (Kilner et al. 1999; Boonstra 2009). Secondly, for the hold period, our experiment naturally requires the recruitment of multiple muscles of the UL to perform the task. Involuntary, muscle-specific interactions during co-activation between muscles of the UL result in lower beta band IMC levels (Lee et al. 2014), which are considered cortical in origin. The same effect may mean that coordinated activation of muscles across the UL may suppress CMC. Similarly, CMC increases to significant levels while executing precise grips involving intrinsic and extrinsic muscles of the hand (Kilner et al. 1999, 2000; Boonstra 2009)the, our experiment used a force target range that required less precision. A low precision task will also result in a decrease of CMC on the beta range (Kristeva-Feige et al. 2002). Additionally, the performance of a cognitive task while executing a motor task also leads to a decrease of CMC (Kristeva-Feige et al. 2002). Our task required a certain amount of cognitive involvement resulting from the spatial nature of the force target; participants had to pay attention to several details on the screen to match the target, which could lead to a decrease of CMC. Finally, force level does not seem to modulate CMC (Mima et al. 1999) on low to moderate contractions, whereas high force levels shift the observed CMC from a beta to a low gamma rhythm (Brown et al. 1998). In our experiment, a force target was set at 40% of MVF for shoulder external rotation, being the weakest direction for force development. Forces generated in all other directions required even lower forces relative to their own MVF. Tasks involving low-to-moderate forces do not modulate coherence (Poston et al. 2010). Our result of no CMC is consistent with this previous finding. Further investigation into the task constraints which permit CMC is warranted.

### Muscle contributions within a synergy

Our results showed higher IMC levels across clusters when comparing IMC between two muscles of high contribution (high–high) than between muscles with high–low contributions within the same synergy. Muscles with a higher contribution within a synergy have roles as the primary movers of the motor task in the synergy’s PD. This has relevance to the integration of theories of common drive and motor control supervision. On one hand, coherence reflects common excitatory drive originating from a neural structure (Danna-Dos-Santos et al. 2014). At the same time, primary movers in a motor task are closely supervised by the motor control system (Krishnamoorthy et al. 2003; Danna-dos-Santos et al. 2007). Therefore, our data imply a link between the level of coherence across muscles and the level of supervision of those muscles by the motor control system. By supporting the existence of common neural drive and closer supervision of the main contributors within each synergy, the implication from our data is that synergies arise from a deliberate neurophysiological control strategy.

Muscle synergies have been analysed from a functional communication point of view by the use of coherence analysis (Marchis et al. 2015). Under dynamic pedalling conditions, only one task-specific synergy, out of four total identified synergies, had significant IMC in the gamma band between high weight muscles (Marchis et al. 2015). Gamma band activity suggests a common cortical input under dynamic conditions (Omlor et al. 2007) such as pedalling. Our results have shown consistent IMC in the alpha band, in every synergy cluster, across topographically scattered muscles. Expression of coherence is sensitive to the experimental conditions and these are therefore the likely cause of the differences between our results and those of other studies. The use of a task where a single synergy is recruited (exerting an isometric for on the PD) on the one hand will constrain the task but, on the other hand, may enhance the underlying recruitment and synchronicity level. The use of a synergy PD may enhance the possibility of identifying underlying neural mechanisms by diminishing the ‘noise’ from multiple synergies being recruited at the same time. In addition, IMC in the alpha band emerges during a sustained low-to-moderate force exertion (Boonstra 2009) paradigm that closely matches our experimental setup. Overall, the exertion of an isometric force towards the synergy PD may explain the consistent level of IMC across all extracted synergies and account for the differences between our results and previous studies.

## Conclusion

The higher IMC of muscles, those with a higher weight within a synergy, are consistent with the idea of synergies being a part of the functional strategy of the CNS to control and build movement. The alpha band IMC suggests the existence of a subcortical mechanism for the generation of muscle synergies. The differential level of IMC between contributors within a synergy may reflect the level of regulation required to achieve the motor task. The existence of a shared input onto synergistic muscles within a synergy supports the idea of neurally-derived muscle synergies, by which a controller reduces the redundant degrees of freedom by a common neural input to synergistic muscles. The lack of CMC is consistent with a task requiring activation of multiple muscles, a low precision force target, and high cognitive involvement.

## Acknowledgements

The authors acknowledge the assistance provided by April Ren and the support provided by the University of Auckland Centre for eResearch. Thanks to MedTech CoRE NZ for the funding provided for the study.

